# Characterization of DSPE-mPEG raw materials from different vendors reveals differences in impurity profiles and polymer chain length polydispersity

**DOI:** 10.1101/2024.01.03.574088

**Authors:** Sven Hackbusch, Sissi White, Min Du

## Abstract

PEGylated lipids such as 1,2-distearoyl-glycero-3-phosphatidylethanolamine-N-methoxy-polyethyleneglycol-2000 (DSPE-mPEG) are essential components of lipid nanoparticle formulations. They have an impact on the efficacy and quality of the drug products they used in, and as such should be carefully characterized. The present work used liquid chromatography with charged aerosol detection and high resolution mass spectrometry to investigate the polydispersity and purity of several lots of DSPE-mPEG, revealing significant differences in the level and identity of impurities. Additionally, mass spectral data showed that there were variations in the polyethylene glycol chain length between lots of comparable purity, which could not be detected by charged aerosol detection alone. The obtained results may be of importance in the development of new lipid nanoparticle based drugs.

**Highlights:**

- Impurity identification of PEGylated lipid raw materials based on MS data
- Determination of PEG polydispersity using high resolution mass spectrometry
- Use of CAD with inverse gradient for impurity quantification

## 1. Introduction

Due to the clinical success in COVID-19 vaccine formulations, lipid nanoparticles (LNPs) have moved to the forefront as drug delivery vehicles for mRNA therapies and others.^[1-4]^ The quality and efficacy of these therapies can be directly impacted by the LNP formulation and as such, its attributes must be monitored carefully.^[5]^ Examples of this for N-oxide impurities have been reported in the literature.^[6]^

While many types of lipids have been explored for LNP formulations, they are typically made up of a combination of four lipid classes: (1) a cationic or ionizable lipid, beneficial for delivery efficacy, (2) a phospholipid, to increase stability of the nanoparticle, (3) cholesterol or derivatives thereof, which increase particle stability and biodistribution, and (4) a polyethylene glycol (PEG)-functionalized lipid, which can modulate particle size and increase stability by decreasing particle aggregation and increasing circulation time.^[1,3-4]^

The structure of the PEGylated lipid has been reported to impact cellular uptake of LNPs, placing an importance on adequate control of molecular weight and polydispersity.^[5,7]^ Additionally, both the impact of PEG chain and lipid anchor length on immunogenicity and pharmacokinetics, respectively, have been reviewed previously.^[8,9]^ However the detailed impurity profiling and polydispersity analysis of PEGylated lipid raw materials has not been previously reported in the literature.

Here, the combination of charged aerosol detection and high-resolution accurate mass (HRAM) MS was used for the raw material characterization of DSPE-mPEG, a PEGylated derivative of 1,2-distearoyl-glycero-3-phosphatidylethanolamine, revealing differences in the impurity profiles and PEG chain length distribution across suppliers.

## 2. Materials and Methods

Samples of 1,2-distearoyl-glycero-3-phosphatidylethanolamine-N-methoxy-polyethyleneglycol-2000 (DSPE-mPEG) were obtained from four different vendors (labelled A-D hereafter) at specified purities ranging from 90 – 99+%. All samples were obtained as powders and separately dissolved in absolute ethanol (Molecular Biology grade, Thermo Scientific), with standard solutions prepared at 1 mg/mL for injection.

Samples were analyzed by reversed phase on a LC/CAD/MS system using a Thermo Scientific Vanquish Inverse Gradient UHPLC system (Germering, Germany) with a Vanquish Charged Aerosol Detector (CAD) and a Thermo Scientific Orbitrap Exploris 120 mass spectrometer (San Jose, CA, USA). Separations were performed with 1 μL injection volumes on a Thermo Scientific™ Hypersil™ GOLD C8 column (1.9 μm, 2.1 × 50 mm) and using a tertiary gradient of (A) 5 mM ammonium formate in water, (B) 5 mM ammonium formate in isopropanol-methanol (70:30, v/v), and (C) isopropanol, at a flow rate of 0.5 mL/min, starting at 0 min with 60:10:30 (%A/%B/%C), increasing to 20:30:50 over 4 min min, changed to 10:40:50 over 2 min, kept constant for 1 min, then returned to initial conditions within 0.1 min and re-equilibrating for 4.9 min. Solvents and additives used were LC-MS grade in all cases.

An inverse gradient with a delay time of 0.58 min was provided by the second pump to maintain a constant mobile phase composition at the CAD.^[10]^ The merged flow from the analytical and inverse gradient pumps was split between the charged aerosol detector and mass spectrometer using a metal T-piece. Mass spectral data were acquired using single polarity Top 4 data-dependent MS^2^ (ddMS^2^) experiments. Complete LC/MS method parameters are detailed in the Supplemental Material. Qualitative and quantitative analysis of the CAD trace data was carried out in Thermo Scientific Chromeleon 7.3.2 CDS and Thermo Scientific™ Compound Discoverer™ 3.3 SP2 software was used for MS compound detection and correlation to CAD peaks, as well as compound identification. Deconvolution of multiply charged species was carried out using the Xtract algorithm in Freestyle 1.8 SP2 software. Detailed data processing parameters are described in the Supplemental Material.

## 3. Results and Discussion

### 3.1 Overview of DSPE-mPEG profiling

As detailed above, PEGylated lipids serve an important role in the LNP formulation. However, from an analytical development standpoint, this class of lipids creates unique challenges, in part due to the variability in the PEG chain length. The polydispersity of the PEG moiety, and its average molecular weight can potentially vary between different vendors and lots.

To investigate this, different lots of DSPE-mPEG were obtained and analyzed using a LC/CAD/MS method described above. As shown in Figure 1 below, the analysis of the different DSPE-mPEG samples indicated the presence of several impurities exceeding the identification threshold of 0.1% cited by ICH guidelines based on the CAD trace.^[11,12]^ Additionally, the main component appeared to have slightly different retention times for the four vendors’ samples.

**Figure 1.**
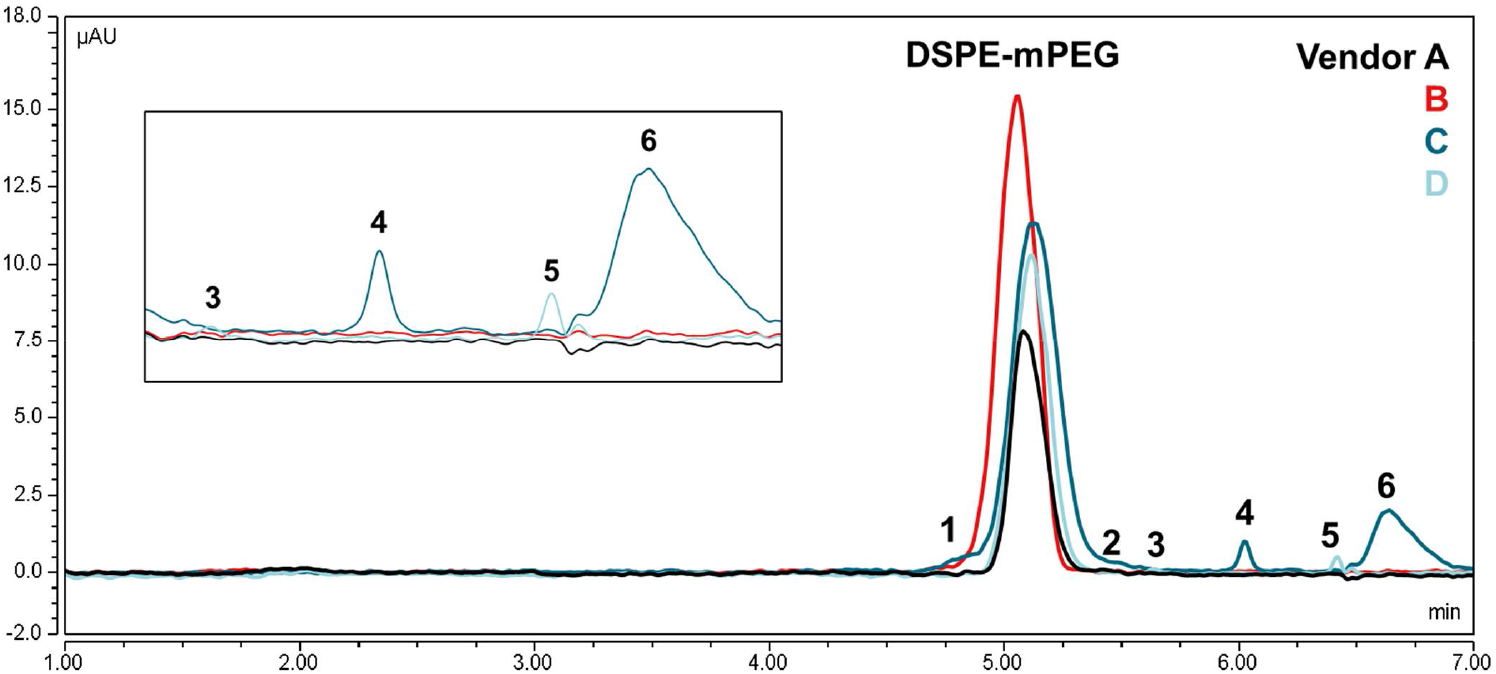
Overlay of the background-subtracted CAD traces for the DSPE-mPEG raw material samples from vendors A-D, showing the apparent retention time shifts between vendors, as well as several impurities, labeled 1-6.

### 3.2 Observed differences in polydispersity

Using the HRAM MS data, it could be determined that this shift in retention times was a result of different polydispersity of the PEG-subunits. Using the Xtract algorithm, the multiply charged ammonium adducts of the DSPE-mPEG peaks at 5.0 min could be deconvoluted to generate the monoisotopic mass distributions for the four samples, which are compared in Figure 2.

**Figure 2.**
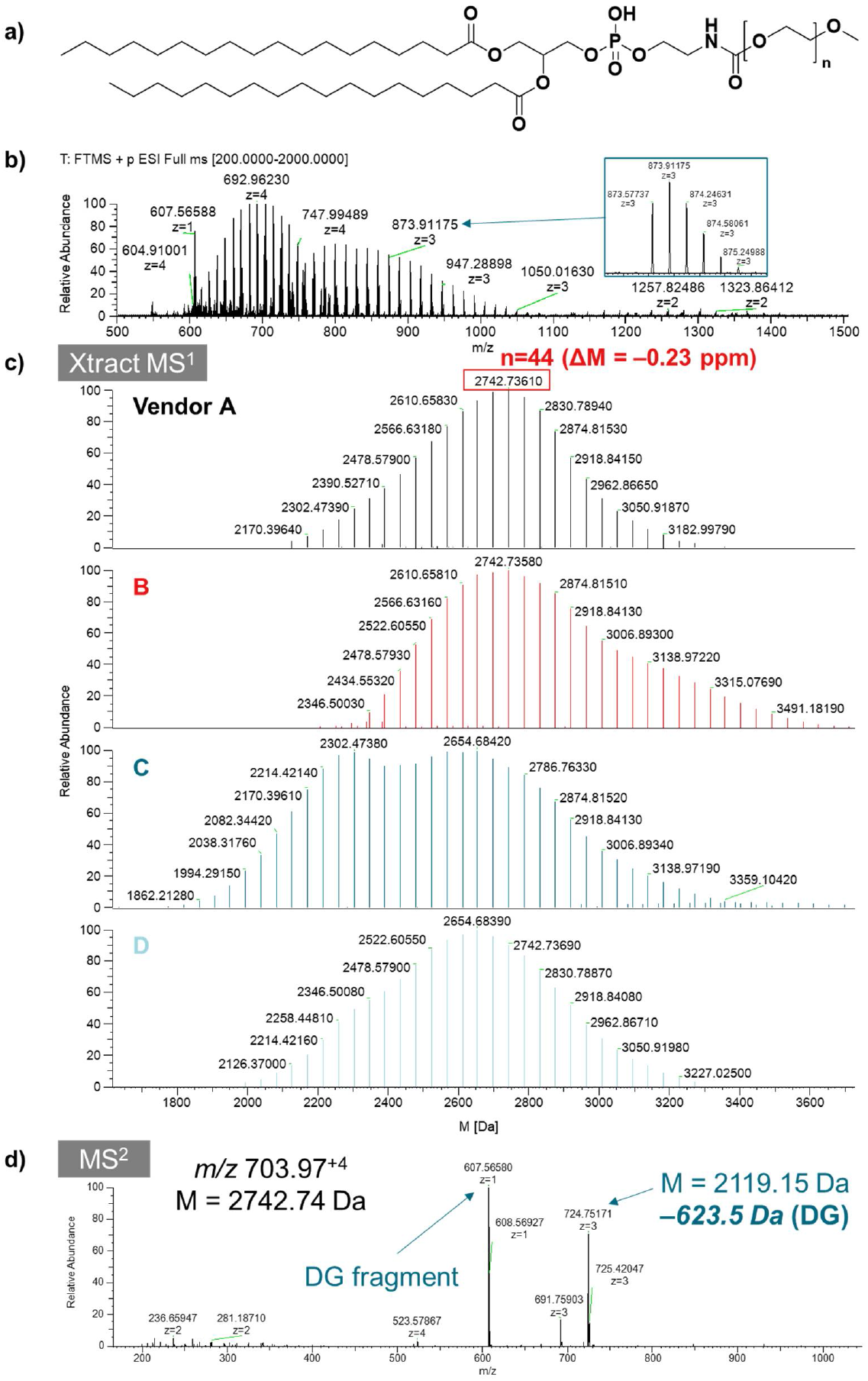
a) Chemical structure of DSPE-mPEG, b) averaged mass spectrum of DSPE-mPEG from 4.8-5.4 min for vendor D, c) comparison of the deconvoluted mass spectra for the four vendors’ materials showing the different polydispersity of the PEG chain lengths, and d) evidence of the distearoyl glyceride in the fragmentation spectrum of the +4 ion of the most abundant species at 2742.74 Da.

The deconvoluted mass spectra showed that the material from vendor B had a distribution with higher masses, in correspondence with its slightly earlier apparent elution, as the higher number of PEG units decreased the compounds’ retention to the C18 column. Interestingly, the distribution for vendor C was found to be bimodal, with two local maxima in terms of PEG chain lengths.

Based on the obtained data, the polydispersity index (PDI) of the different raw materials could be calculated as follows from the weight average molar mass (M_w_) and number average molar mass (M_n_), with the results summarized in Table 1:

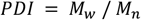

**Table 1.** Calculated numerical and weight average molar masses and PDI for DSPE-mPEG from the different vendors based on the data presented in Figure 2b.

| Vendor | $M_n$ | $M_w$ | PDI |
| --- | --- | --- | --- |
| A | 2688.535 | 2706.022 | 1.006504 |
| B | 2834.391 | 2859.173 | 1.008743 |
| C | 2543.749 | 2581.398 | 1.014801 |
| D | 2627.195 | 2650.282 | 1.008788 |

Notably, even though the calculated PDIs for the lots from vendor B and D were quite similar, their average molar masses were markedly different, which may impact the LNP properties if one were substituted for the other.

### 3.3 Impurity identification

To determine the identity of the impurities detected in the CAD data of the different DSPE-mPEG materials highlighted in Figure 1, both automatic processing using the Compound Discoverer software for non-PEGylated impurities and manual interrogation of multiply charged PEGylated impurities was carried out, as detailed hereafter. A summary of the impurity profiling results for DSPE-mPEG is given in Table 2.

**Table 2.**
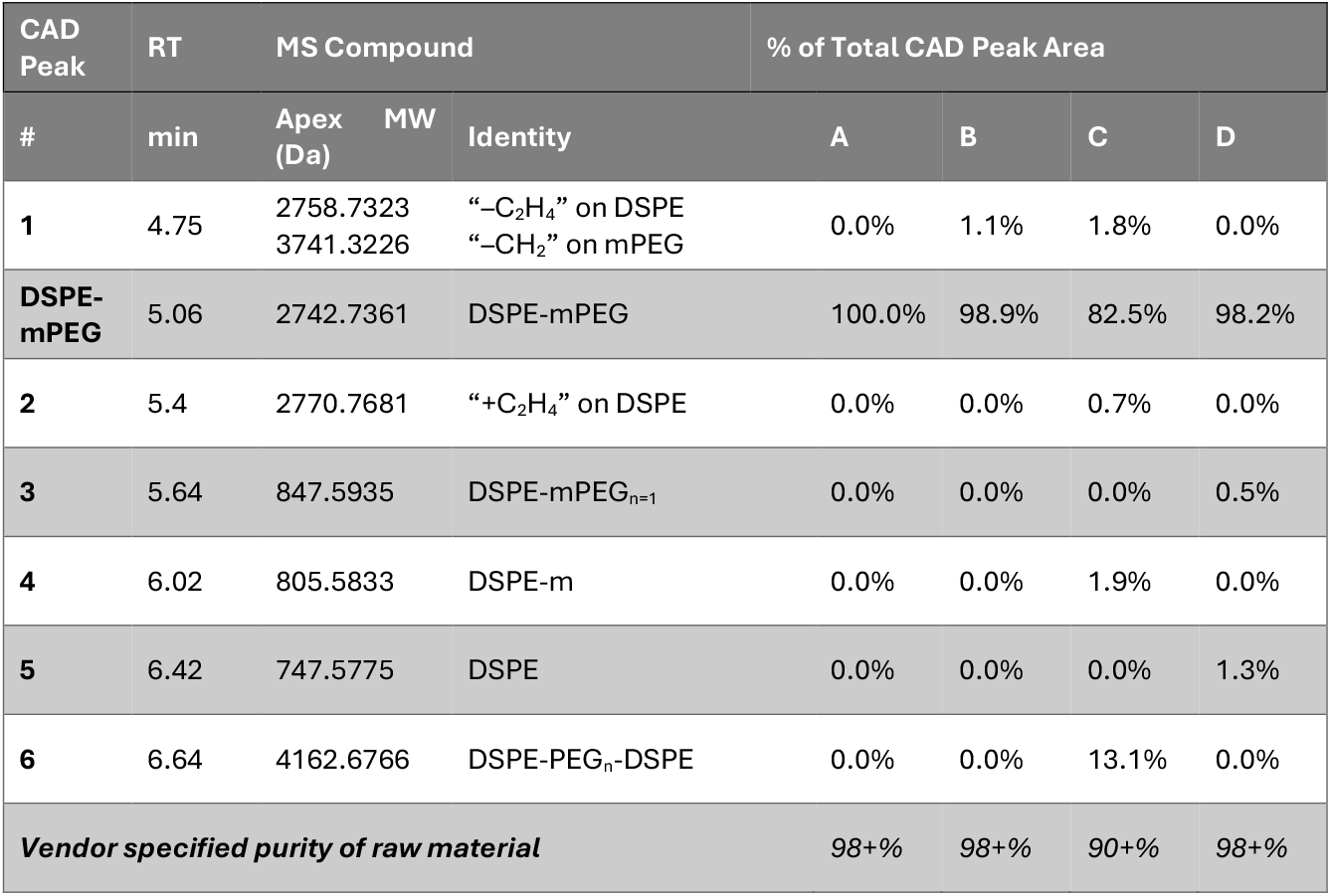
Summary of the impurity profiling results for the DSPE-mPEG samples from vendors A-D.

Impurities **3, 4** and **5** were found not to contain a PEG moiety, since only singly charged components below *m/z* 1000 were detected for them in the MS data.

Impurity **5**, which was detected in material from vendor D, could be identified as 1,2-distearoyl-glycero-3-phosphatidylethanolamine (DSPE), with the characteristic headgroup loss of 141 Da leading to the formation of the distearoyl glycerol ion at *m/z* 607 seen in the MS^2^ spectrum (Figure S2 in the Supplemental Material).

The MS compounds detected for impurities **3** and **4** differed by +100.0160 Da and +58.0058 Da, relative to impurity **5**. Based on the high-resolution accurate mass data, these could be confidently determined to result from an elemental composition difference of +C_4_H_4_O_3_ and +C_2_H_2_O_2_ relative to the composition of C_41_H_82_NO_8_P for DSPE, respectively.

Based on this, and the presence of the characteristic *m/z* 607 ion in their MS^2^ spectra, the structures shown in Figure 3 were proposed as their respective identities.

**Figure 3.**
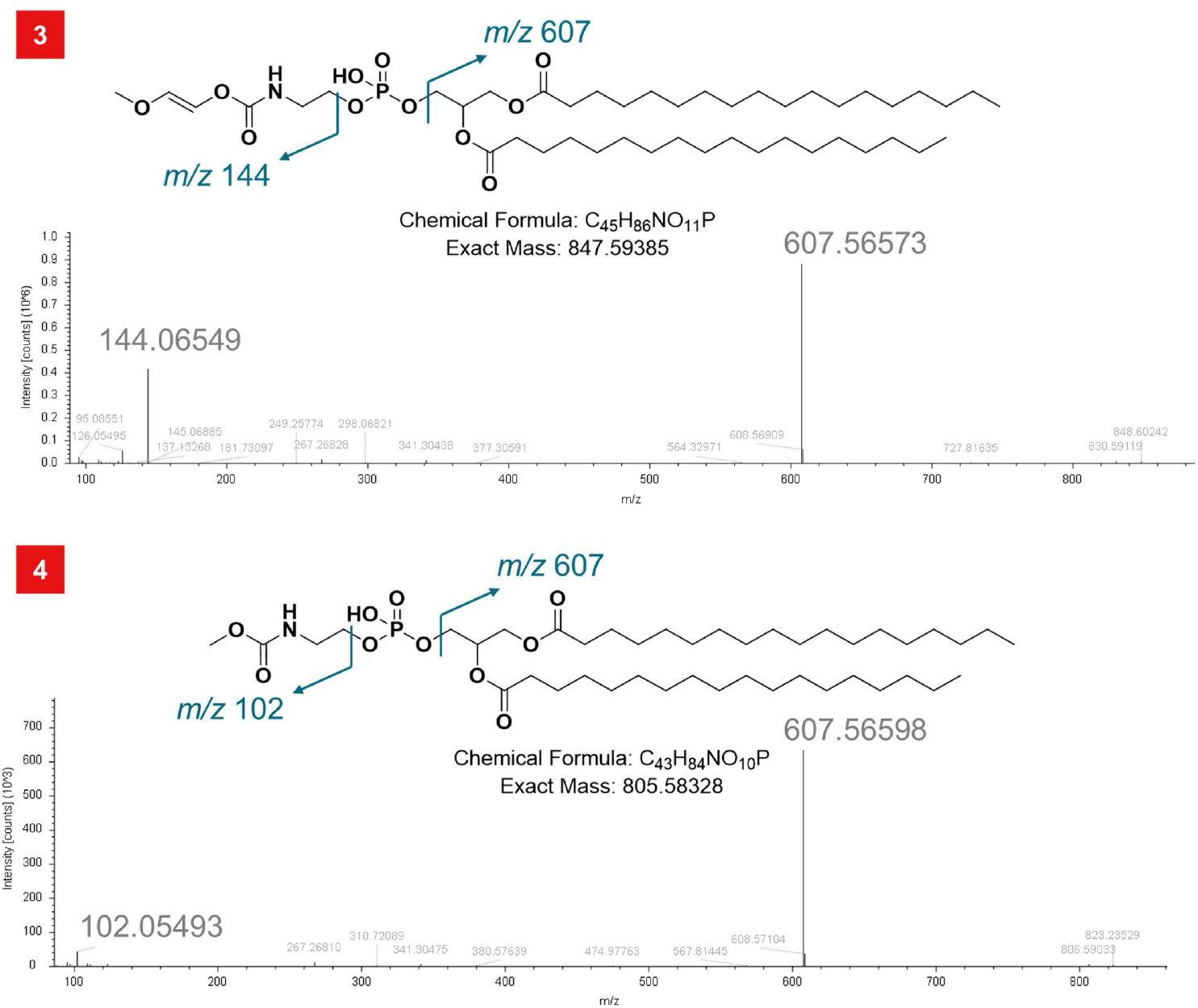
MS^2^ spectra and proposed chemical structures for impurities 3 and 4 of DSPE-mPEG.

**Figure 4.**
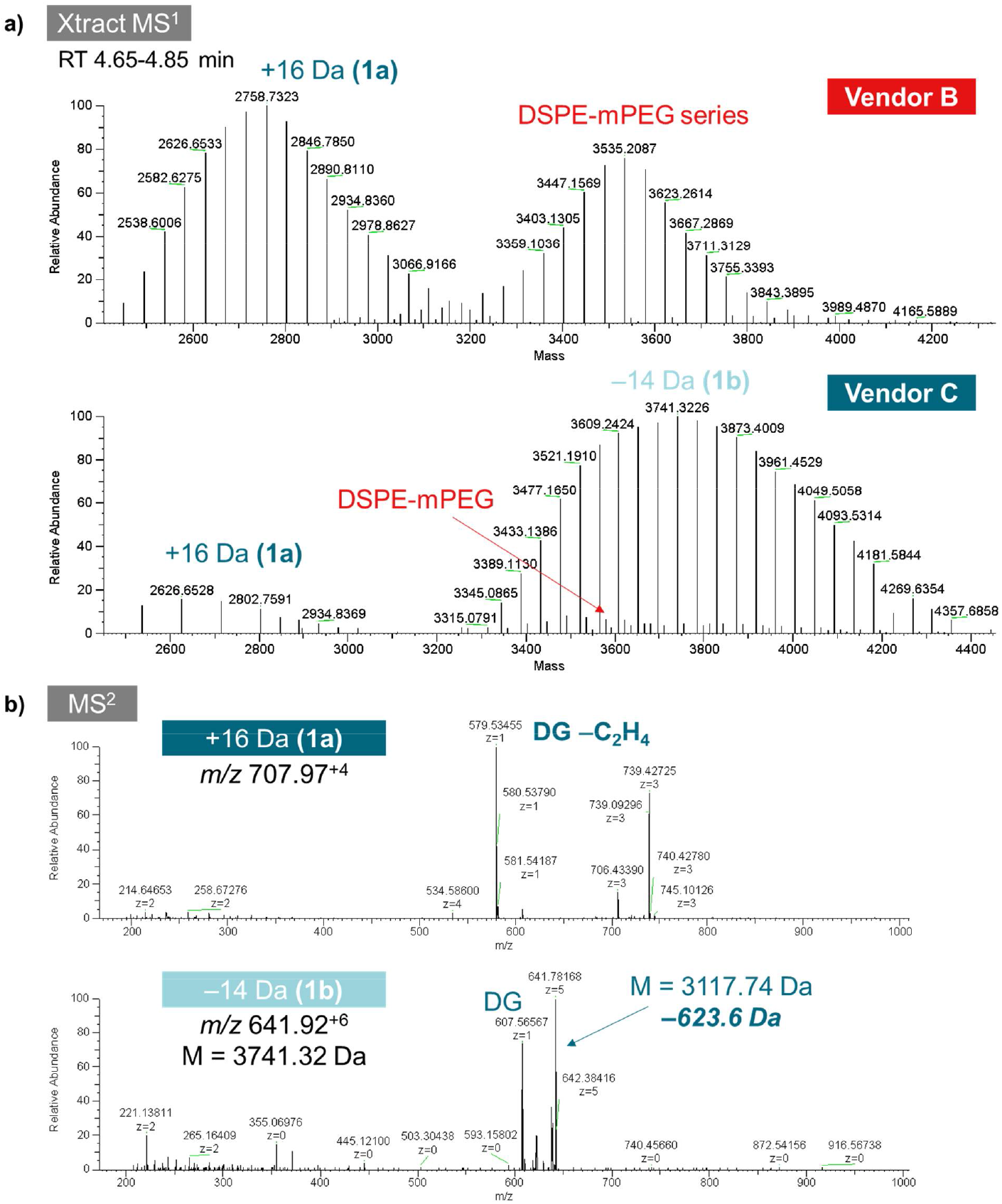
a) Deconvolved average mass spectra for RT 4.65-4.85 min of materials B and C, indicating the presence of different PEGylated impurities with mass differences of +16 Da (1a) and –14 Da (1b) relative to the parent compound series of DSPE-mPEG, respectively. B) MS^2^ fragmentation spectra of corresponding precursors for impurities 1a and 1b, indicating a “–C_2_H_4_” modification of the distearoyl moiety in 1a corresponding to a mass shift of –28 Da and no modification on the distearoyl moiety in 1b.

Both compounds could reasonably be formed as side products in the methylation of unPEGylated DSPE intermediates in the synthesis of DSPE-mPEG.

Meanwhile, impurities **1, 2** and **6** were found to display the polydispersity characteristics of the PEG moiety. They were interrogated after deconvolution of the MS data at the respective elution time ranges, as detailed hereafter.

As shown in Figure 7, the deconvolution of the average mass spectrum from 4.65-4.85 min for impurity **1** of materials B and C revealed two distinct PEGylated species differing from the parent structure by +16 Da (**1a**) and –14 Da (**1b**), respectively, along with higher *n* species of the parent DSPE-mPEG_n_.

Investigation of corresponding MS^2^ spectra for the most abundant ion of **1a** provided evidence for a modification of the distearoyl glycerol moiety for **1a**, corresponding to “–C_2_H_4_” (–28 Da), and resulting in the observed fragment mass of *m/z* 579 – meaning the ion with apparent +16 Da mass difference contained one additional PEG unit (+44 Da) but two fewer carbons on the diglyceride. Meanwhile, for **1b**, the distearoyl glycerol fragment was detected unchanged at *m/z* 607, meaning that the observed –14 Da modification was the result of a “–CH_2_” transformation on the mPEG moiety, possibly from the absence of the terminal methyl group.

Analysis of impurity peak **2** indicated the presence of a PEGylated series of compounds with a mass shift of –16 Da relative to the parent structure, which could be determined to result from an addition of “+C_2_H_4_” to the DSPE moiety, as evidenced by a shift in the observed fragment ion, in analogy to **1a** (Figure S3). In both cases, the likely origin of these impurities was proposed to be respective impurities in the DSPE starting material with different fatty acid chain lengths.

Lastly, the average mass spectrum across the elution time of impurity **6** could be deconvolved to reveal a PEGylated compound series with a significantly increased apex mass, shifted +1419.94 Da relative to the apex mass of the parent species in material from vendor A. This large mass shift pointed to the presence of a higher *n* of PEG units, while the fragmentation spectrum indicated the presence of a second DSPE moiety. Taken together, these observed modifications could be explained by a dimeric structure formed in the reaction of PEG with DSPE units at both ends (Figure S4). The observed monoisotopic mass of 4162.6766 Da matched the theoretical mass for the proposed structure with a mass error of only 1.25 ppm, increasing the confidence in this putative identification.

## 4. Conclusions

Here, we present the profiling and impurity identification of a PEGylated lipid raw material. The analysis of different lots of DSPE-mPEG raw material revealed significant differences in their molecular weight distribution and impurity profiles. While the PDIs of the investigated lots were generally comparable, the average molecular weight was found to differ even for lots with similar PDI. Secondly, multiple impurities with and without the polyethylene glycol moiety could be detected and assigned structures based on the mass spectral data, likely resulting from starting materials (**5**), side products (**4**), as well as structural analogs of DSPE-mPEG created from impurities in the starting materials used in its synthesis (**1a**). These findings emphasize the need for in-depth characterization of raw materials and the utility of mass spectral data for the analysis of PEGylated lipids.

## Conflict of Interest section

S.H., S.W. and M.D. are employees of Thermo Fisher Scientific, whose instruments and software were used in this research.

## Author contributions

**S.H**.: Conceptualization, Methodology Investigation, Data curation, Formal Analysis, Visualization, Writing – Original Draft; **S**.**W**.: Conceptualization, Methodology, Writing – Review & Editing; **M**.**D**.: Conceptualization, Resources, Writing – Review & Editing.

