## Supplemental Information for "Characterization of DSPE-mPEG raw materials from different vendors reveals differences in impurity profiles and polymer chain length polydispersity"

for

### Table S1. UHPLC method parameters

| **Parameter** | **Value** |
| --- | --- |
| Column | Thermo Scientific™ Hypersil™ GOLD C8, 1.9 µm, 2.1 × 50 mm (P/N 25202-052130) |
| Mobile Phases | A: 5 mM Ammonium formate in 100% H_2_O  B: 5 mM Ammonium formate in 70% IPA/30% MeOH  C: 100% IPA |
| Flow Rate | 0.5 mL/min |
| Column Temperature | 50 °C (still air mode) |
| Autosampler Temperature | 6 °C |
| Injection Volume | 1 µL |
| Needle Wash | 5 mM Ammonium formate in 70% IPA/30% MeOH, before and after draw |
| Divert valve timing | Flow to waste from 0 – 1 min |
| CAD Settings | Evaporation Temperature 35 °C, Power Function 1.00, Data Collection Rate 5 Hz, Filter 3.6 |

### Table S2. UHPLC Gradient conditions

| **Time (min)**  (**for inverse gradient, add 0.58 min) | **Analytical Gradient Pump** | | | **Inverse Gradient Pump** | | |
| --- | --- | --- | --- | --- | --- | --- |
|  | **%A** | **%B** | **%C** | **%A** | **%B** | **%C** |
| 0 | 60 | 10 | 30 | 0 | 30 | 70 |
| 4 | 20 | 30 | 50 | 30 | 10 | 60 |
| 6 | 10 | 40 | 50 | 50 | 0 | 50 |
| 7 | 10 | 40 | 50 | 50 | 0 | 50 |
| 7.1 | 60 | 10 | 30 | 0 | 30 | 70 |
| 12 | 60 | 10 | 30 | 0 | 30 | 70 |

### Table S3. MS source conditions

| **Parameter** | **Value** |
| --- | --- |
| Spray Voltage | + 3250 V / – 3000 V |
| Sprayer Position | 1.5, M/H, center |
| Vaporizer temperature | 300 °C |
| Ion transfer tube temperature | 350 °C |
| Sheath gas | 50 a.u. |
| Aux gas | 10 a.u. |
| Sweep Gas | 1 a.u. |

### Table S4. MS Experiment FullScan - ddMS^2^ method parameters

| **Parameter** | **Value** |
| --- | --- |
| MS^1^ Resolution | 120,000 @ *m/z* 200 |
| MS^1^ Mass Range | *m/z* 200-2000 |
| RF Level, % | 70 |
| Easy-IC | Scan-to-Scan |
| MS^2^ Isolation Window (*m/z*) | 1.5 |
| HCD Collision Energies (Normalized, %) | 10,20,30 |
| MS^2^ Resolution | 30,000 @ *m/z* 200 |
| Maximum Injection Time (ms) | Auto |
| Intensity Threshold | 1.0e5 |
| Dynamic Exclusion | Custom, 8 s, Exclude isotopes, 5 ppm mass tolerance |

### Compound Discoverer processing workflow

Created with Discoverer version: 3.3.1.111, *non-default parameters denoted*

[Input Files (0)]

-->Create Analog Trace (33)

-->Select Spectra (46)

*Lower RT Limit: 1 min*

*Upper RT Limit : 7 min*

[Select Spectra (46)]

-->Align Retention Times (42)

[Align Retention Times (42)]

-->Detect Compounds (64)

*Min. Peak Intensity: 1000000*

*Ions: [M+2NH4]+2, [M+3NH4]+3, [M+4NH4]+4, [M+5NH4]+5,	[M+6NH4]+6, [M+H]+1, [M+Na]+1, [M+NH4]+1*

-->Find Expected Compounds (65)

[Detect Compounds (64)]

-->Group Compounds (28)

*Peak Rating Threshold: 4, Number of Files: 1*

-->Merge Features (14)

[Group Compounds (28)]

-->Fill Gaps (62)

-->Calculate Mass Defect (67)

-->Assign Compound Annotations (30)

-->Predict Compositions (45)

*Max. Element Counts: C190 H390 Br3 Cl4 N10 O100 S5*

[Generate Expected Compounds (35)]

-->Find Expected Compounds (65)

*- Compounds: DSPE-PEG2000-n37 (C117 H232 N O47 P)*

*- Apply Dealkylation: True*

*- Transformations: - Phase I: (not specified) - Phase II: (not specified) - Others: Demethylation (C H3 -> H), FA Chain lengthening ( -> C2 H4), FA Chain shortening (C2 H4 -> ), PEG subunit ( -> C2 H4 O) - Max. # Phase II: 1 - Max. # All Steps: 3*

[Find Expected Compounds (65)]

-->Group Expected Compounds (37)

*Peak Rating Threshold: 4, Number of Files: 1*

-->Merge Features (14)

[Fill Gaps (62)]

-->Mark Background Compounds (44)

[Assign Compound Annotations (30)]

-->Generate Molecular Networks (69)

[Group Expected Compounds (37)]

-->Mark Background Compounds (63)

[Create Analog Trace (33)]

[Mark Background Compounds (44)]

[Calculate Mass Defect (67)]

[Generate Molecular Networks (69)]

[Predict Compositions (45)]

[Merge Features (14)]

[Mark Background Compounds (63)]

[Differential Analysis (61)]

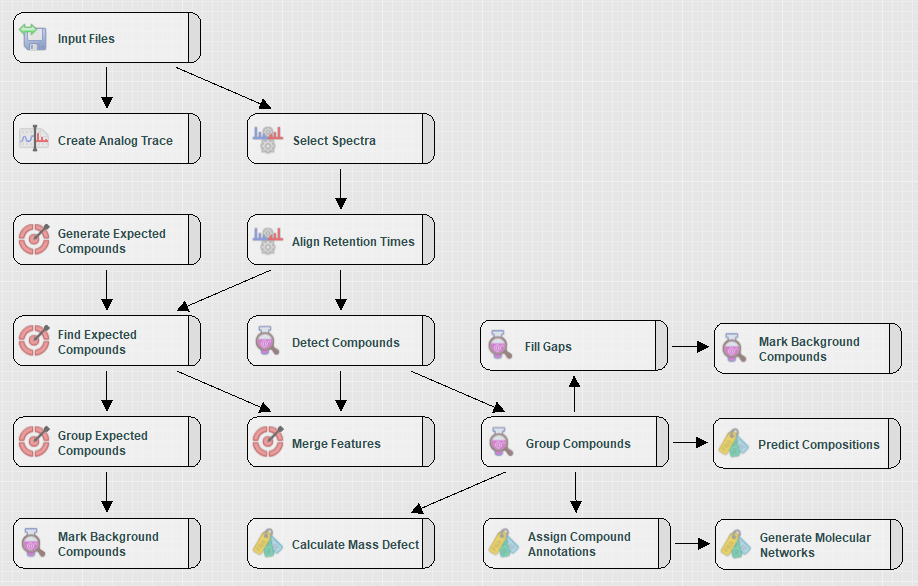

**Figure S1. Workflow tree in Compound Discoverer 3.3**

### Table S5. Freestyle 1.8 SP2 Xtract Deconvolution Parameters

| **Parameter** | **Value** |
| --- | --- |
| Output Mass | M |
| Adduct Element | Custom, 18.03383 |
| Charge Range | 2 to 7 |
| Analyzer Type | OT |
| Rel. Abund. Threshold (%) | 0 |
| Isotope Table | Protein |
| Negative Charge | Unchecked |
| Min Num Detected Charge | 2 |

### Table S6. Deconvolved monoisotopic mass and peak intensity data for DSPE-mPEG from vendors A-D used in the PDI calculations shown in Table 1

(data from averaged mass spectra 4.8-5.4 min)

| **A** | | **B** | | **C** | | **D** | |
| --- | --- | --- | --- | --- | --- | --- | --- |
| **Mono-isotopic mass** | **Intensity** | **Mono-isotopic mass** | **Intensity** | **Mono-isotopic mass** | **Intensity** | **Mono-isotopic mass** | **Intensity** |
| 2346.5003 | 1.37E+07 | 1994.292 | 3.67E+06 | 1774.16 | 9.42E+05 | 2126.37 | 4.84E+06 |
| 2390.5273 | 2.79E+07 | 2038.318 | 6.58E+06 | 1818.186 | 1.99E+06 | 2170.396 | 7.96E+06 |
| 2434.5532 | 4.79E+07 | 2082.344 | 1.06E+07 | 1862.213 | 4.32E+06 | 2214.422 | 1.25E+07 |
| 2478.5793 | 7.03E+07 | 2126.37 | 1.66E+07 | 1906.239 | 8.36E+06 | 2258.447 | 1.91E+07 |
| 2522.6055 | 9.16E+07 | 2170.396 | 2.46E+07 | 1950.266 | 1.47E+07 | 2302.474 | 2.69E+07 |
| 2566.6316 | 1.09E+08 | 2214.422 | 3.53E+07 | 1994.292 | 2.36E+07 | 2346.5 | 3.32E+07 |
| 2610.6581 | 1.21E+08 | 2258.448 | 4.95E+07 | 2038.318 | 3.38E+07 | 2390.527 | 3.99E+07 |
| 2654.6838 | 1.29E+08 | 2302.474 | 5.94E+07 | 2082.344 | 4.75E+07 | 2434.553 | 4.89E+07 |
| 2698.71 | 1.31E+08 | 2346.501 | 6.61E+07 | 2126.37 | 6.14E+07 | 2478.579 | 6.02E+07 |
| 2742.7358 | 1.32E+08 | 2390.527 | 7.23E+07 | 2170.396 | 7.58E+07 | 2522.606 | 7.11E+07 |
| 2786.7624 | 1.28E+08 | 2434.553 | 8.12E+07 | 2214.421 | 8.88E+07 | 2566.632 | 8.09E+07 |
| 2830.7891 | 1.22E+08 | 2478.579 | 9.18E+07 | 2258.448 | 9.75E+07 | 2610.658 | 9.15E+07 |
| 2874.8151 | 1.13E+08 | 2522.606 | 1.03E+08 | 2302.474 | 9.95E+07 | 2654.684 | 9.84E+07 |
| 2918.8413 | 1.01E+08 | 2566.631 | 1.11E+08 | 2346.501 | 9.51E+07 | 2698.71 | 1.04E+08 |
| 2962.8663 | 8.58E+07 | 2610.658 | 1.16E+08 | 2390.527 | 9.04E+07 | 2742.736 | 1.05E+08 |
| 3006.893 | 7.37E+07 | 2654.684 | 1.18E+08 | 2434.553 | 9.08E+07 | 2786.763 | 1.00E+08 |
| 3050.9186 | 6.54E+07 | 2698.71 | 1.14E+08 | 2478.579 | 9.19E+07 | 2830.789 | 9.10E+07 |
| 3094.9458 | 6.01E+07 | 2742.737 | 1.07E+08 | 2522.606 | 9.68E+07 | 2874.815 | 7.78E+07 |
| 3138.9722 | 5.47E+07 | 2786.763 | 9.94E+07 | 2566.632 | 1.00E+08 | 2918.842 | 5.99E+07 |
| 3182.998 | 4.99E+07 | 2830.789 | 8.75E+07 | 2610.658 | 9.92E+07 | 2962.867 | 4.59E+07 |
| 3227.0249 | 4.42E+07 | 2874.815 | 7.51E+07 | 2654.684 | 1.00E+08 | 3006.893 | 3.32E+07 |
| 3271.051 | 3.86E+07 | 2918.841 | 6.22E+07 | 2698.71 | 9.53E+07 | 3050.919 | 2.45E+07 |
| 3315.0769 | 3.30E+07 | 2962.867 | 4.75E+07 | 2742.737 | 9.01E+07 | 3094.946 | 1.85E+07 |
| 3359.1026 | 2.65E+07 | 3006.893 | 3.67E+07 | 2786.763 | 8.47E+07 | 3138.972 | 1.23E+07 |
| 3403.1295 | 2.10E+07 | 3050.92 | 2.80E+07 | 2830.789 | 7.64E+07 | 3182.998 | 8.29E+06 |
| 3447.1554 | 1.62E+07 | 3094.946 | 2.15E+07 | 2874.815 | 6.77E+07 | 3227.025 | 4.79E+06 |
| 3491.1819 | 1.19E+07 | 3138.972 | 1.61E+07 | 2918.841 | 5.61E+07 | 3271.05 | 2.77E+06 |
| 3535.208 | 7.79E+06 | 3182.999 | 1.10E+07 | 2962.867 | 4.58E+07 | 3359.104 | 8.06E+05 |
| 3579.2333 | 5.08E+06 | 3227.025 | 7.10E+06 | 3006.893 | 3.65E+07 |  |  |
| 3623.2599 | 2.86E+06 | 3271.051 | 4.29E+06 | 3050.92 | 3.12E+07 |  |  |
| 3667.2836 | 1.60E+06 | 3359.103 | 1.09E+06 | 3094.946 | 2.52E+07 |  |  |
| 3711.3121 | 7.18E+05 |  |  | 3138.972 | 2.08E+07 |  |  |
|  |  |  |  | 3182.999 | 1.64E+07 |  |  |
|  |  |  |  | 3227.025 | 1.24E+07 |  |  |
|  |  |  |  | 3271.051 | 9.04E+06 |  |  |
|  |  |  |  | 3315.077 | 6.73E+06 |  |  |
|  |  |  |  | 3359.104 | 4.35E+06 |  |  |
|  |  |  |  | 3403.13 | 2.93E+06 |  |  |
|  |  |  |  | 3447.16 | 1.99E+06 |  |  |
|  |  |  |  | 3491.183 | 1.31E+06 |  |  |

Figure S2. Identification of impurity 5 **based on correlation of the CAD trace and the XIC for a MS compound with MW 747.5775, which could be identified as 1,2-distearoyl-glycero-3- phosphatidylethanolamine (DSPE) from its fragmentation spectrum, with its structure given below.**

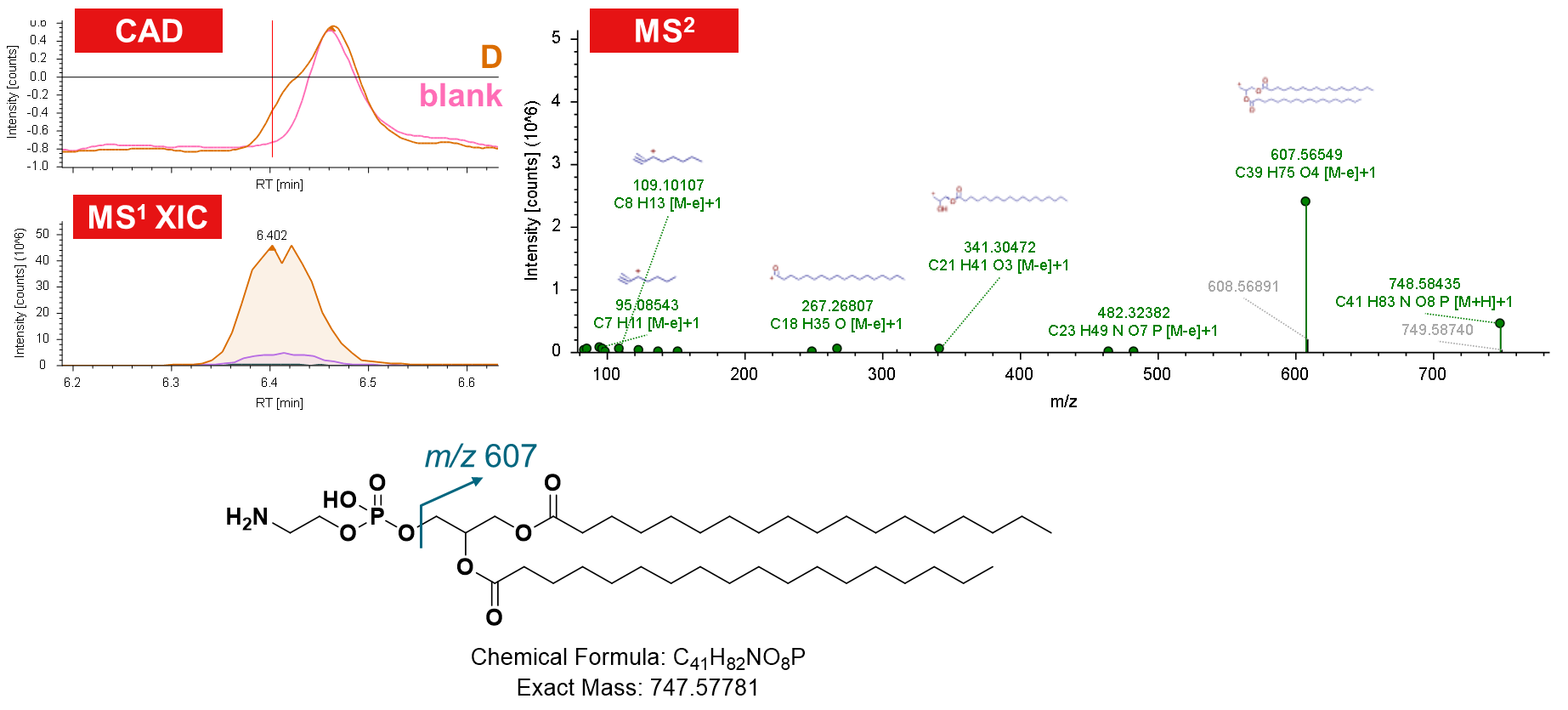

Figure S3. Identification of impurity 2 **based on a) deconvolved average mass spectrum for RT 5.3-5.5 min of material C, indicating the presence of PEGylated impurities with a mass difference of –16 Da relative to the parent compound series of DSPE-mPEG, part of which was also seen in the spectrum, and b) MS^2^ fragmentation spectrum of a corresponding precursor for impurity 2, indicating a “+C_2_H_4_” modification of the distearoyl moiety corresponding to a mass shift of +28 Da.**

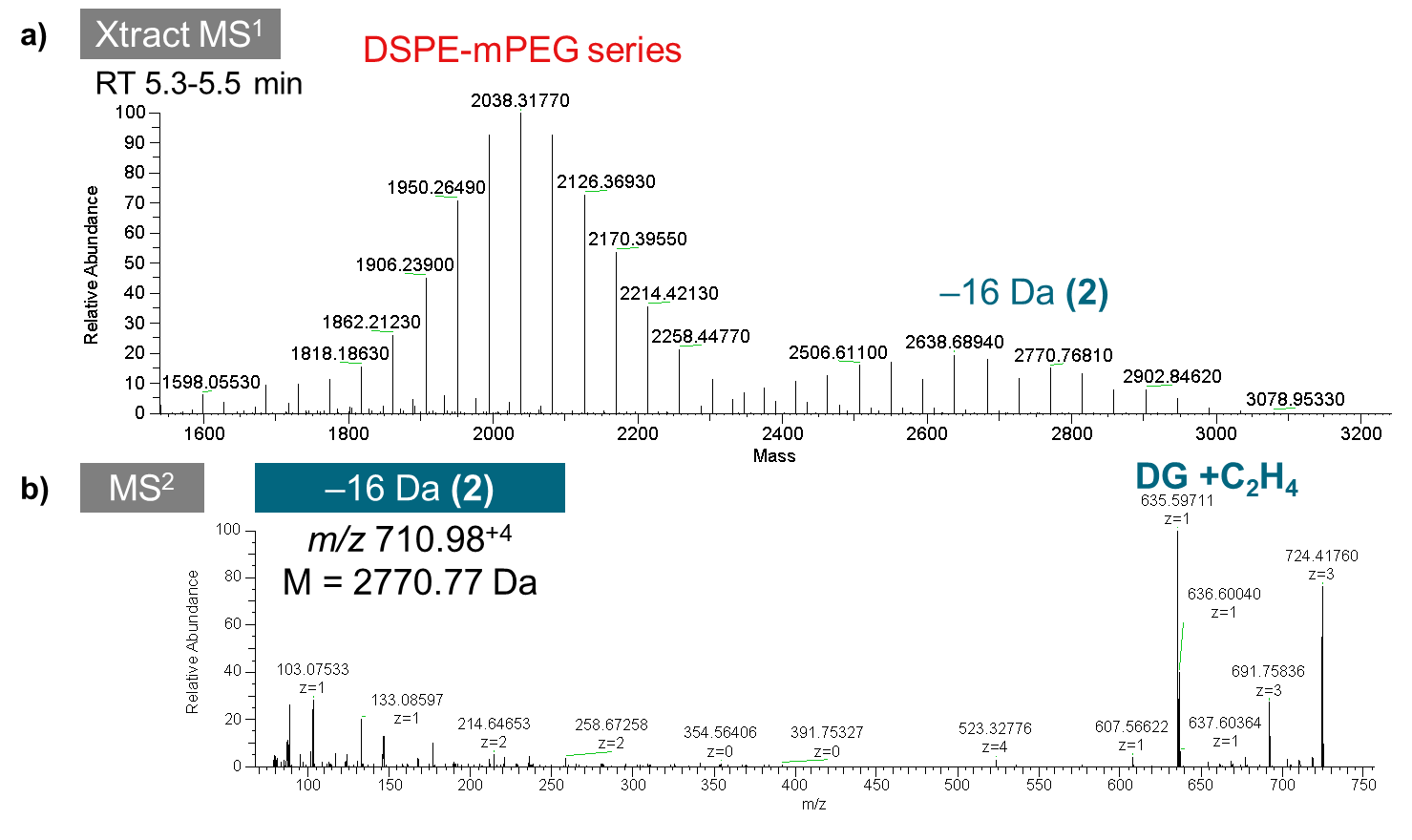

Figure S4. Identification of Impurity 6 **based on a) the deconvoluted mass spectrum for impurity 6 (RT 5.44-5.84 min) and b) representative fragmentation spectrum for the +5 ion of the most abundant species (4162.7 Da), providing evidence for two losses of a distearoyl glycerol moiety, which could be explained by the dimeric structure shown in c).**

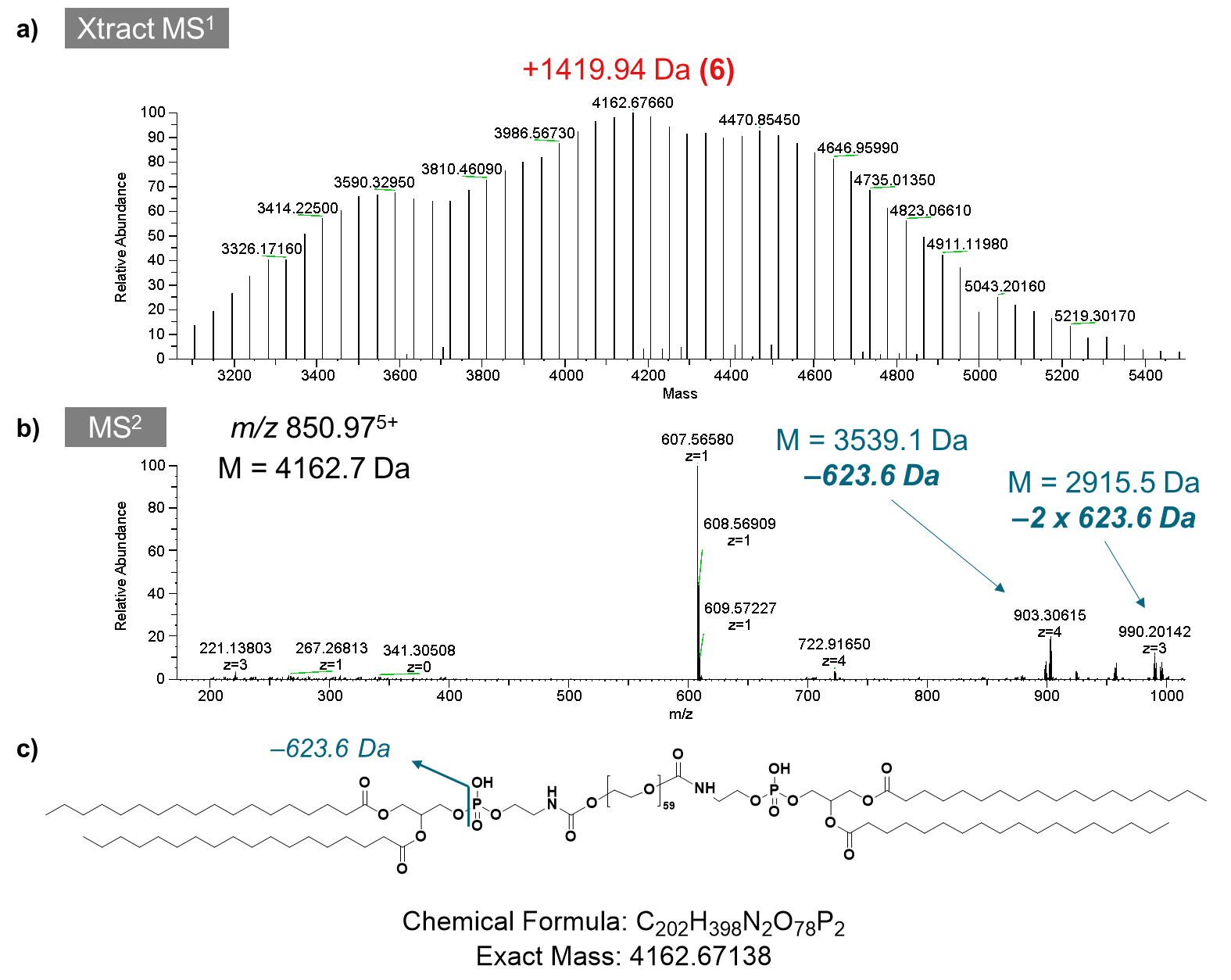
